# MethylGPT: a foundation model for the DNA methylome

**DOI:** 10.1101/2024.10.30.621013

**Authors:** Kejun Ying, Jinyeop Song, Haotian Cui, Yikun Zhang, Siyuan Li, Xingyu Chen, Hanna Liu, Alec Eames, Daniel L McCartney, Riccardo E. Marioni, Jesse R. Poganik, Mahdi Moqri, Bo Wang, Vadim N. Gladyshev

## Abstract

DNA methylation serves as a powerful biomarker for disease diagnosis and biological age assessment. However, current analytical approaches often rely on linear models that cannot capture the complex, context-dependent nature of methylation regulation. Here we present MethylGPT, a transformer-based foundation model trained on 226,555 (154,063 after QC and deduplication) human methylation profiles spanning diverse tissue types from 5,281 datasets, curated 49,156 CpG sites, and 7.6 billion training tokens. MethylGPT learns biologically meaningful representations of CpG sites, capturing both local genomic context and higher-order chromosomal features without external supervision. The model demonstrates robust methylation value prediction (Pearson R=0.929) and maintains stable performance in downstream tasks with up to 70% missing data. Applied to age prediction across multiple tissue types, MethylGPT achieves superior accuracy compared to existing methods. Analysis of the model’s attention patterns reveals distinct methylation signatures between young and old samples, with differential enrichment of developmental and aging-associated pathways. When finetuned to mortality and disease prediction across 60 major conditions using 18,859 samples from Generation Scotland, MethylGPT achieves robust predictive performance and enables systematic evaluation of intervention effects on disease risks, demonstrating potential for clinical applications. Our results demonstrate that transformer architectures can effectively model DNA methylation patterns while preserving biological interpretability, suggesting broad utility for epigenetic analysis and clinical applications.

## Introduction

DNA methylation is an epigenetic modification where methyl groups are added to cytosine residues at CpG dinucleotides. This modification regulates gene expression by recruiting methyl-CpG binding proteins and modifying chromatin accessibility ^1^. DNA methylation regulates multiple biological processes through distinct mechanisms. During development, dynamic methylation changes guide cellular differentiation by silencing lineage-inappropriate genes and activating cell-type-specific programs ^2^. Methylation also maintains genomic stability through the repression of transposable elements ^3^.

Beyond its fundamental role in gene regulation, DNA methylation exhibits key characteristics of an ideal biomarker: stability in the resting state, but with dynamic response to environmental factors, accessibility in various biological specimens, and early alterations preceding clinical manifestations ^4^. Genome-wide methylation profiling has revealed distinctive signatures across numerous pathological states, enabling molecular diagnostics, particularly in cancer detection and cardiovascular risk assessment ^5^.

Alongside disease prediction, age-associated methylation patterns also enable the development of highly accurate “epigenetic aging clocks” ^6^. These clocks have evolved from simple age predictors to sophisticated biomarkers of biological aging, with recent advances such as DunedinPACE ^7^, GrimAge ^8^, causality-enriched clocks ^9^, and the high-dimensional ageome ^10^, demonstrating strong associations with health outcomes and mortality risk. Notably, these methylation-based aging indices often outperform conventional clinical measures in predicting age-related diseases and longevity ^11,12^, highlighting their potential for monitoring therapeutic interventions targeting the aging process.

However, several analytical challenges impede the clinical implementation of methylation-based diagnostics. Current computational approaches predominantly rely on linear models and simple statistical methods, which are fundamentally limited in their ability to capture complex, non-linear relationships in methylation data. These linear models assume independence between CpG sites, failing to account for the regulatory networks and higher-order interactions that characterize methylation patterns. Moreover, the same DNA methylation pattern may have different biological implications depending on the cellular and tissue context: a complexity that linear models are unable to capture ^13–15^. The limitations of linear models become even more apparent when dealing with technical artifacts, including batch effects and missing data, which introduce substantial non-linear variability in methylation measurements ^16^. The field urgently needs a unified analytical framework capable of modeling complex, non-linear patterns, accounting for contextdependent effects, and performing robust pattern analysis across diverse clinical contexts.

Recent advances in artificial intelligence, particularly transformer architectures and foundation models ^17^, have revolutionized the analysis of complex biological sequences. Foundation models have emerged across multiple omics layers: for proteomics, ESM-2/ESM-3 ^18,19^ and AlphaFold2/AlphaFold3 ^20,21^ have achieved unprecedented accuracy in structure prediction and function annotation; for genomics, Enformer ^22^ and Evo ^23^ have demonstrated capability in predicting gene regulation and variant effects. In the single-cell domain, models like Geneformer ^24^, scGPT ^25^, and scFoundation ^26^ have enabled zero-shot cell-type classification and in-silico perturbation. And more recently, the Precious3GPT has emerged as a multimodal transformer model integrating multi-omics data for aging research and drug discovery ^27^

These foundation models demonstrate remarkable capability in learning comprehensive biological patterns that generalize across tasks. However, despite the success of foundation models across various omics layers, DNA methylation analysis lacks such a unified approach, relying instead on task-specific models that fail to capture the full complexity of methylation patterns. The achievements of foundation models in related domains suggest that a similar approach could transform methylation analysis by providing a unified framework that preserves biological context while enabling adaptations to diverse specific tasks.

Here, we introduce MethylGPT (Fig. 1a), a transformer-based foundation model for DNA methylation. Trained on methylation profiles from over 150,000 human samples spanning diverse tissue types, MethylGPT implements a novel embedding strategy to capture methylation patterns at physiologically relevant CpG sites. This approach enables unified analysis of DNA methylation data across multiple experimental contexts and downstream applications, including age prediction and disease association detection.

**Figure 1.**
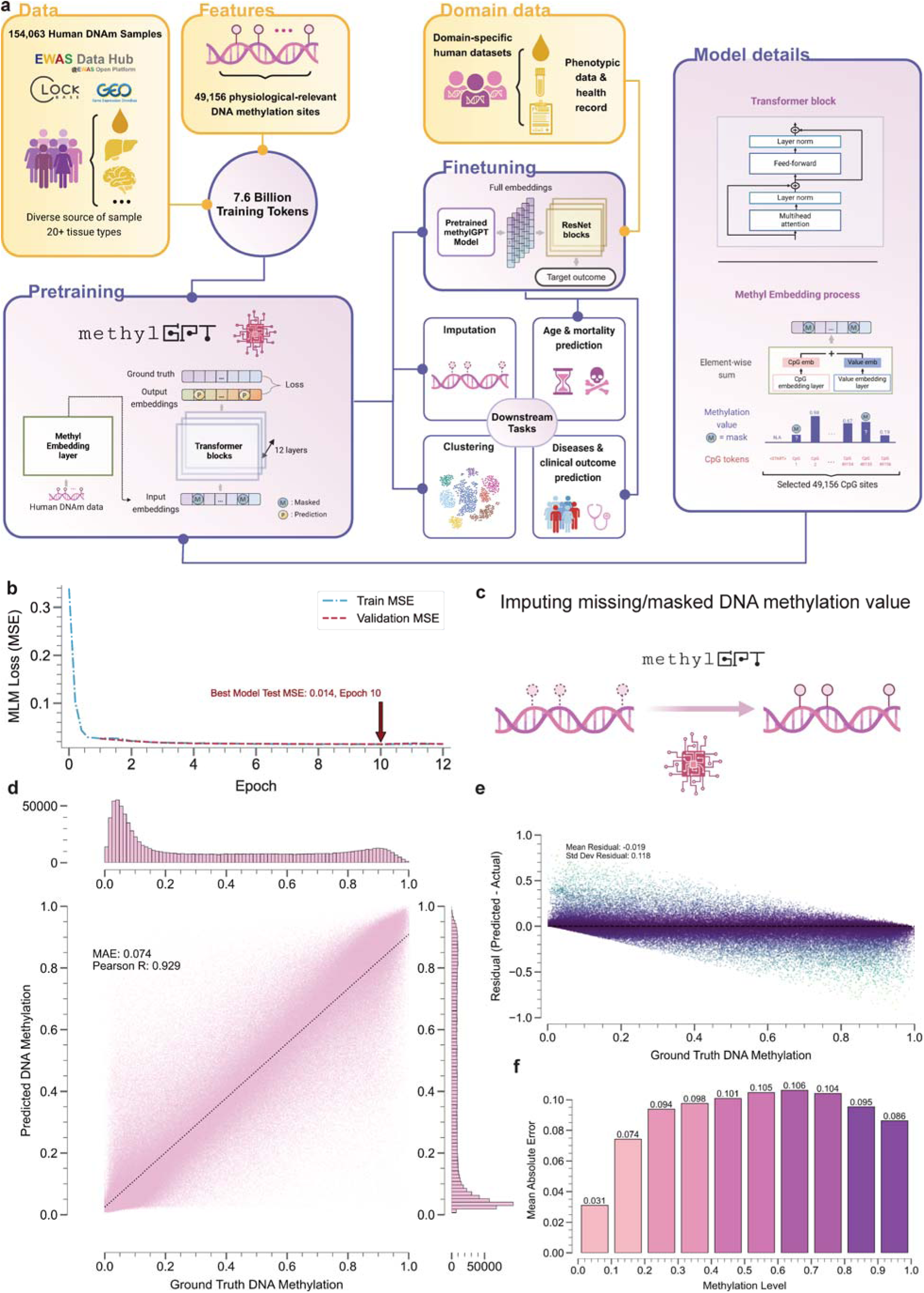
Overview of MethylGPT architecture and performance. **a.** Model architecture diagram showing data flow from 154,063 human DNAm samples through feature extraction (49,156 CpG sites) to generate 7.6 billion training tokens. Components, including transformer block details and the methyl embedding process, are highlighted. **b.** Training curve showing MLM loss over epochs, with train and validation MSE trajectories converging at epoch 10 (Best Model Test MSE: 0.014). **c.** Illustration of the imputing process for missing/masked DNA methylation values using MethylGPT. **d.** Joint density plot showing the correlation between predicted and ground truth DNA methylation values (Pearson R: 0.929, MAE: 0.074). **e.** Residual plot showing prediction errors across different methylation levels. **f.** Bar plot showing mean absolute error across different methylation levels (0.0-1.0).

## Results

### Development and validation of MethylGPT

To enable the pretraining of large-scale model, we collected 226,555 human DNA methylation profiles from 5,281 datasets through the EWAS Data Hub and Clockbase ^28,29^. After quality control and deduplication, we used 154,063 samples to pretrain MethylGPT. The model focuses on 49,156 physiologically-relevant CpG sites, selected based on association with EWAS traits (Methods) ^30^. These methylation profiles, representing samples from over 20 different tissue types, were processed to generate 7.6 billion training tokens (CpG sites), enabling comprehensive coverage of methylation patterns across the human epigenome.

The core architecture of MethylGPT consists of a methylation embedding layer followed by 12 transformer blocks (Fig. 1a). Our methylation embedding process captures both the CpG site tokens and their methylation states through an element-wise attention mechanism. This design enables the model to learn complex dependencies between distant CpG sites while maintaining local methylation context. The model was pre-trained using two complementary loss functions: a masked language modeling (MLM) loss where the model predicts methylation levels for 30% randomly masked CpG sites and a reconstruction loss where the Classify token (CLS) embedding is used to reconstruct the complete DNA methylation profile.

To evaluate the model’s performance, we first assessed its ability to predict DNA methylation values at masked CpG sites in the test set. During training, the model achieved rapid convergence with minimal overfitting, reaching a best model test mean squared error (MSE) of 0.014 at epoch 10 (Fig. 1b). The model demonstrated robust prediction accuracy across different methylation levels, achieving an overall mean absolute error (MAE) of 0.074 and a Pearson correlation coefficient of 0.929 between predicted and actual methylation values (Fig. 1c-f).

### MethylGPT learns biologically meaningful CpG representations

To investigate whether MethylGPT captures biologically relevant DNA methylation features, we analyzed the learned representations of 49K CpG sites in the embedding space (Fig. 2a). Dimensionality reduction using UMAP revealed distinct patterns in the contextualized CpG embedding space (Fig. 2b). CpG sites clustered according to their genomic contexts, with clear separation based on CpG island relationships (island, shore, shelf, and other regions), suggesting that our model learned underlying regulatory features of the methylome without explicit supervision.

**Figure 2.**
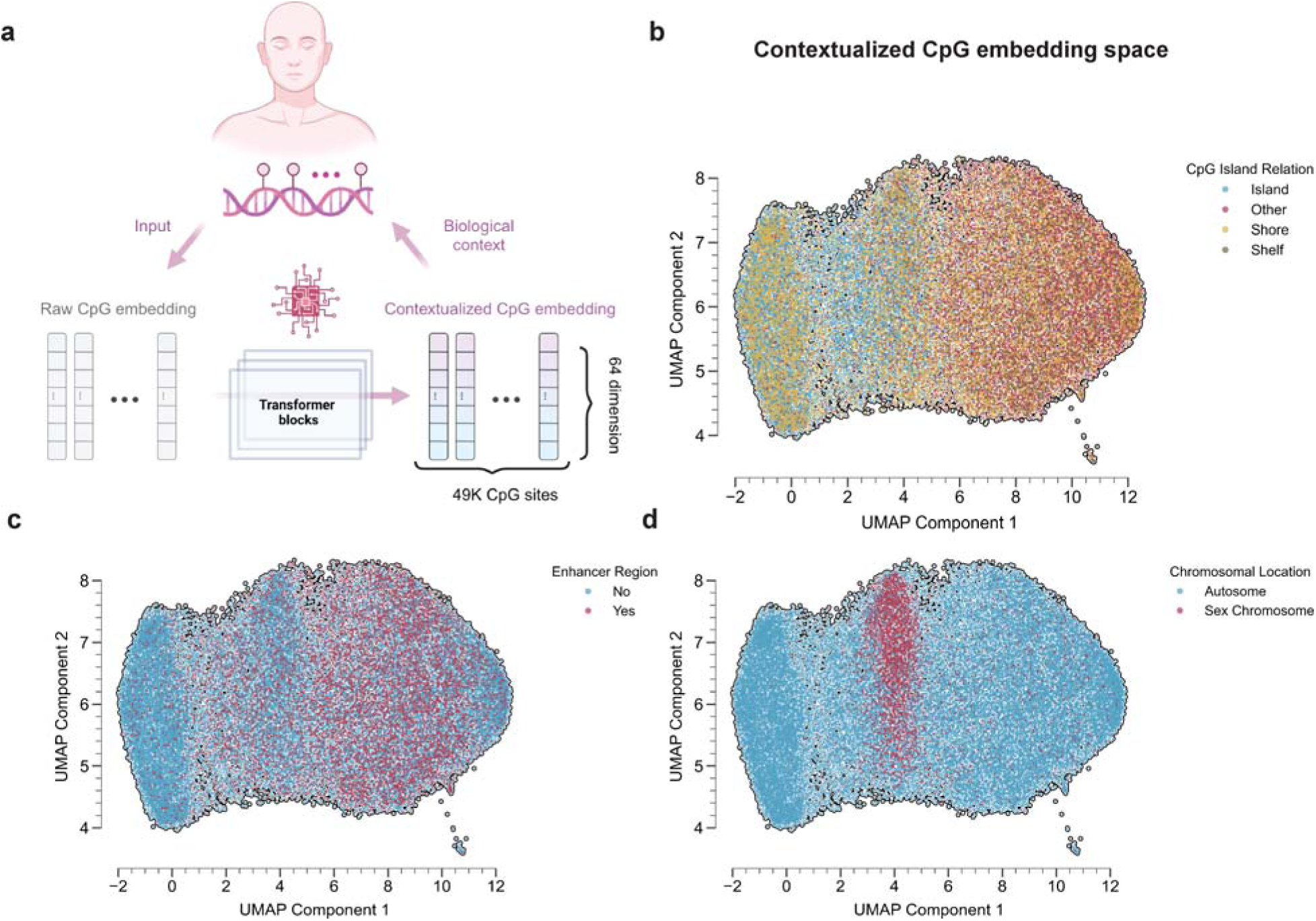
Analysis of contextualized CpG embedding space. **a.** Schematic illustration of the CpG embedding process, showing the transformation from raw CpG input to contextualized embeddings through transformer blocks. **b.** UMAP visualization of 49K CpG sites colored by CpG island relationship (Island, Shore, Shelf, Other). **c.** UMAP plot highlighting enhancer regions (Yes/No) in the embedding space. **d.** UMAP visualization showing the separation of CpG sites by chromosomal location, with distinct clustering of sex chromosomes and autosomes.

The embedding space organization reflected known biological properties of DNA methylation regulation. CpG sites within enhancer regions showed distinct clustering patterns (Fig. 2c), consistent with their specialized regulatory roles. Furthermore, the embeddings demonstrated a clear separation of sex chromosomes from autosomes (Fig. 2d). This organization indicates that MethylGPT successfully captured both local sequence context and higher-order chromosomal features that influence methylation patterns.

The transformer architecture enabled our model to learn these complex relationships through its attention mechanism, which integrates both local CpG site features and broader genomic context (Fig. 2a) instead of treating CpG sites as independent entities as in previous methods.

### MethylGPT learns tissue-specific and sex-specific methylation patterns

To evaluate whether MethylGPT captures biologically meaningful sample-level features, we analyzed the zero-shot embedding spaces of DNA methylation samples before and after model processing. The contextualized sample embeddings from MethylGPT showed clear biological organization, with distinct clustering patterns by tissue type and sex (Fig. 3a). Major tissue types, including whole blood, brain, liver, and skin, formed well-defined clusters, suggesting that MethylGPT successfully learned tissue-specific methylation signatures without explicit supervision. Notably, batch effects were not significant in the observed embeddings (Fig. 3b). MethylGPT embeddings also revealed strong sex-specific methylation patterns across tissues (Fig. 3c). Male and female samples showed consistent separation in the embedding space, reflecting known sex-specific methylation differences.

**Figure 3.**
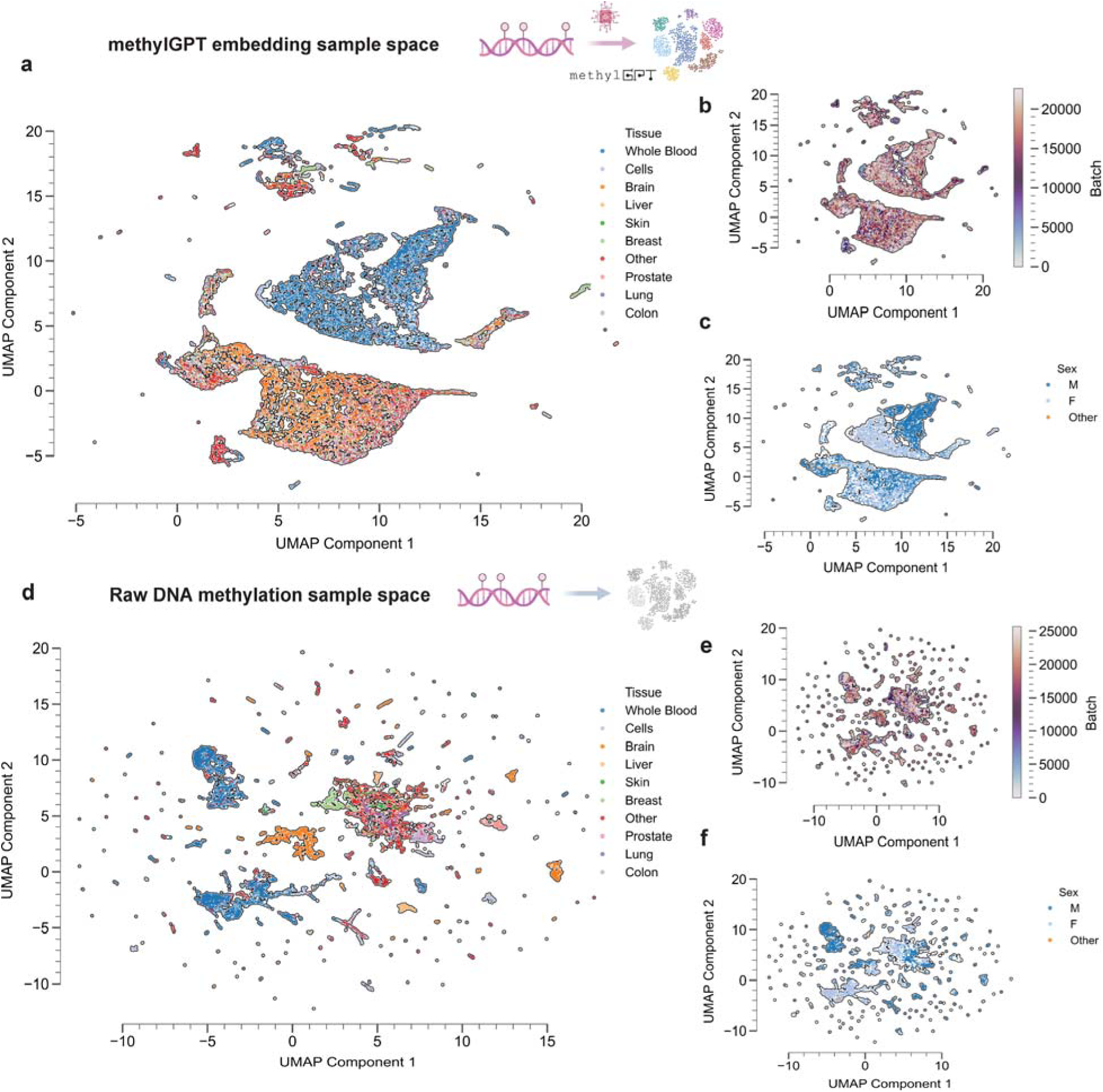
Sample-level embedding analysis. **a.** UMAP visualization of MethylGPT sample embeddings colored by tissue type, showing distinct clustering of major tissue types including whole blood, brain, liver, and skin. **b.** Sample density plot of the embedding space highlighting minimal batch effects. **c.** Sex-specific clustering in the embedding space, displaying a clear separation between male and female samples. **d-f.** Comparative analysis of raw DNA methylation sample embeddings, showing less distinct clustering by tissue type (d), more pronounced batch effects (e), and weaker separation by sex (f).

The superiority of MethylGPT’s learned representations becomes apparent when compared to the raw methylation data directly generated UMAP embeddings (Fig. 3d-f). While raw methylation profiles showed some degree of tissue-specific clustering, the boundaries between different tissue types were less distinct, and the overall organization was more diffuse. The raw data embeddings exhibited less defined tissue-specific clusters (Fig. 3d), stronger batch-specific clustering (Fig. 3e), and weaker sex-specific separation (Fig. 3f), highlighting MethylGPT’s ability to enhance biologically relevant signals through its contextualized embedding approach.

### MethylGPT enables accurate age prediction across diverse tissue types

To evaluate MethylGPT’s capability in downstream applications, we first assessed its performance in predicting chronological age from DNA methylation patterns. We utilized a diverse dataset of 11,453 samples spanning multiple tissue types ^31^, with an age distribution ranging from 0 to 100 years (Fig. 4a). The majority of samples were derived from whole blood (47.2%) and brain tissue (34.5%), providing broad coverage of physiologically distinct methylation patterns.

**Figure 4.**
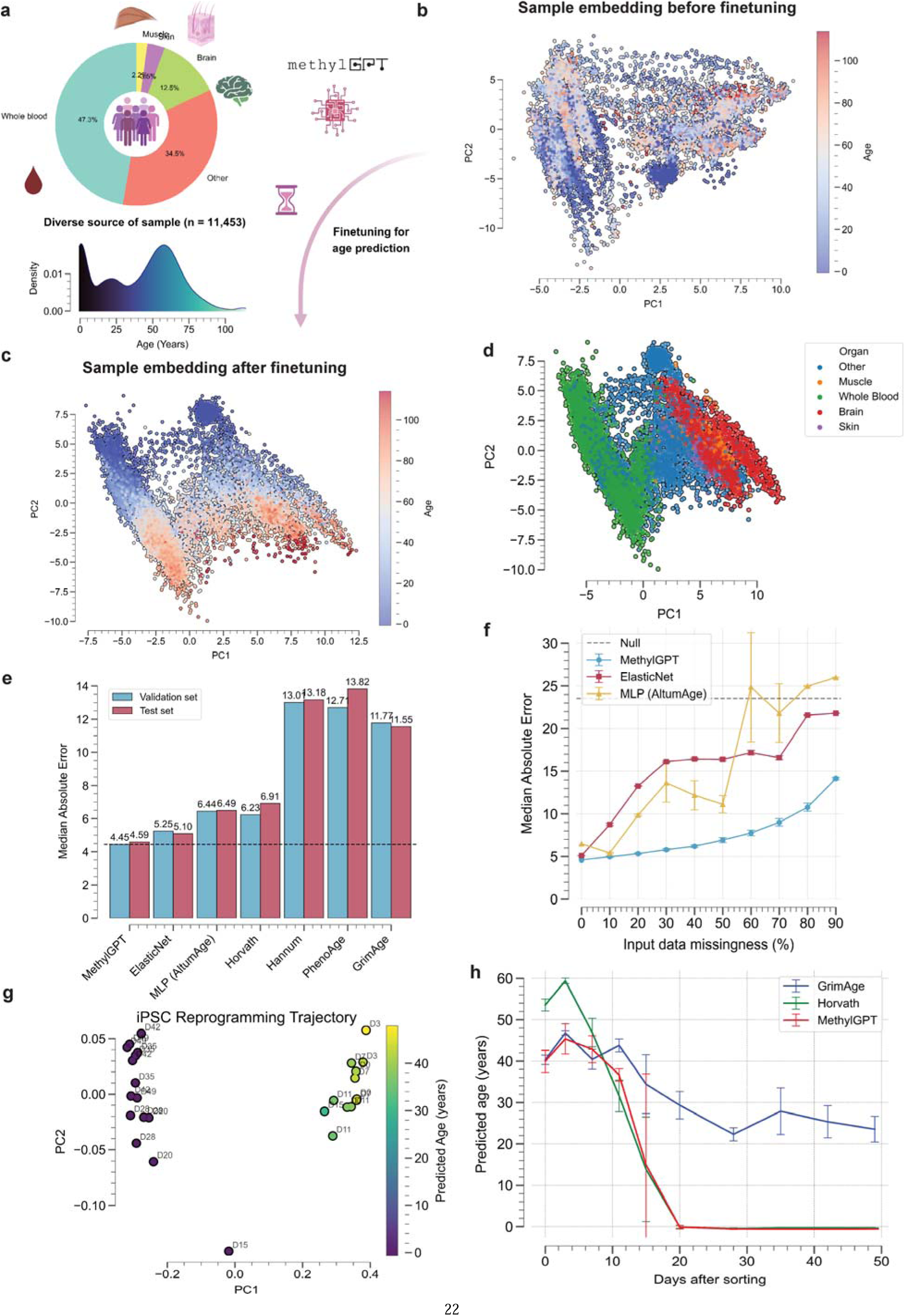
Age prediction performance and robustness analysis. **a.** Sample composition pie chart showing tissue distribution within the age finetuning dataset (n=11,453) and age distribution density plot. **b.** PCA visualization of sample embeddings before fine-tuning, colored by age. **c.** Sample embeddings after fine-tuning for age prediction, showing enhanced age-related organization. **d.** Tissue-specific clustering was maintained after fine-tuning. **e.** Benchmark comparison of age prediction performance across different methods on validation and test datasets. Median Absolute Errors are annotated. **f.** Robustness analysis showing prediction performance under increasing levels of missing data (10-90%) on test dataset for different methods. **g**. Principal component analysis of MethylGPT embeddings during iPSC reprogramming, colored by predicted age, showing progressive trajectory towards younger methylation states. **h**. Comparison of predicted age trajectories during iPSC reprogramming across different epigenetic clocks (GrimAge, Horvath’s clock) and MethylGPT, demonstrating consistent detection of rejuvenation effects. Error bars represent standard deviation across replicate samples.

The pre-trained MethylGPT embeddings showed inherent age-related organization even before fine-tuning (Fig. 4b), suggesting that the model captured age-associated methylation features during pre-training. After fine-tuning for age prediction, the sample embeddings demonstrated stronger age-dependent clustering (Fig. 4c) while maintaining tissue-specific patterns (Fig. 4d).

We compared MethylGPT’s age prediction performance against existing methods, including ElasticNet ^32^, MLP (AltumAge) ^31^, Horvath’s skin and blood clock ^33^, and other established age predictors. MethylGPT achieved superior accuracy with a median absolute error (MedAE) of 4.45 years on the validation set, outperforming other methods (Fig. 4e). This improvement was consistent across both validation and test sets, demonstrating the model’s robust generalization capability.

Notably, MethylGPT showed remarkable resilience to missing data, a common challenge in methylation analysis. We systematically evaluated prediction performance under increasing levels of data missingness (10-90%). MethylGPT maintained stable performance with up to 70% missing data, significantly outperforming both ElasticNet and Multi-Layer Perceptron (MLP) approaches (Fig. 4f). This robustness suggests that the model’s contextualized embeddings effectively capture redundant age-related signals across multiple CpG sites, enabling reliable predictions despite incomplete methylation profiles.

To further validate MethylGPT’s ability to capture biologically meaningful age-related patterns, we analyzed DNA methylation profiles during iPSC reprogramming ^34^. The model’s embeddings revealed a clear rejuvenation trajectory (Fig. 4g), with samples progressively shifting towards a younger methylation state over the reprogramming time course. Notably, when compared with conventional epigenetic clocks (Horvath’s clock and GrimAge), MethylGPT showed consistent detection of rejuvenation effects, predicting a significant decrease in epigenetic age during reprogramming (Fig. 4h). This agreement with established aging biomarkers, while accounting for the broader epigenomic context through the transformer architecture, provides independent support for iPSC reprogramming as a rejuvenation method rather than merely a cell identity transformation. The predicted age trajectory showed a sharp decline after day 20 of reprogramming, reaching near-zero predicted ages by day 30, consistent with the restoration of a pluripotent epigenetic state.

### Age-specific attention patterns reveal distinct methylation signatures

To investigate how MethylGPT processes age-related methylation patterns, we analyzed the model’s multi-head self-attention weights (Fig. 5a). By examining the attention weight matrices, we observed that the model learned distinct patterns of CpG site interactions between young (age < 20) and old (age > 60) samples, suggesting that the transformer architecture captures age-specific relationships in methylation data.

**Figure 5.**
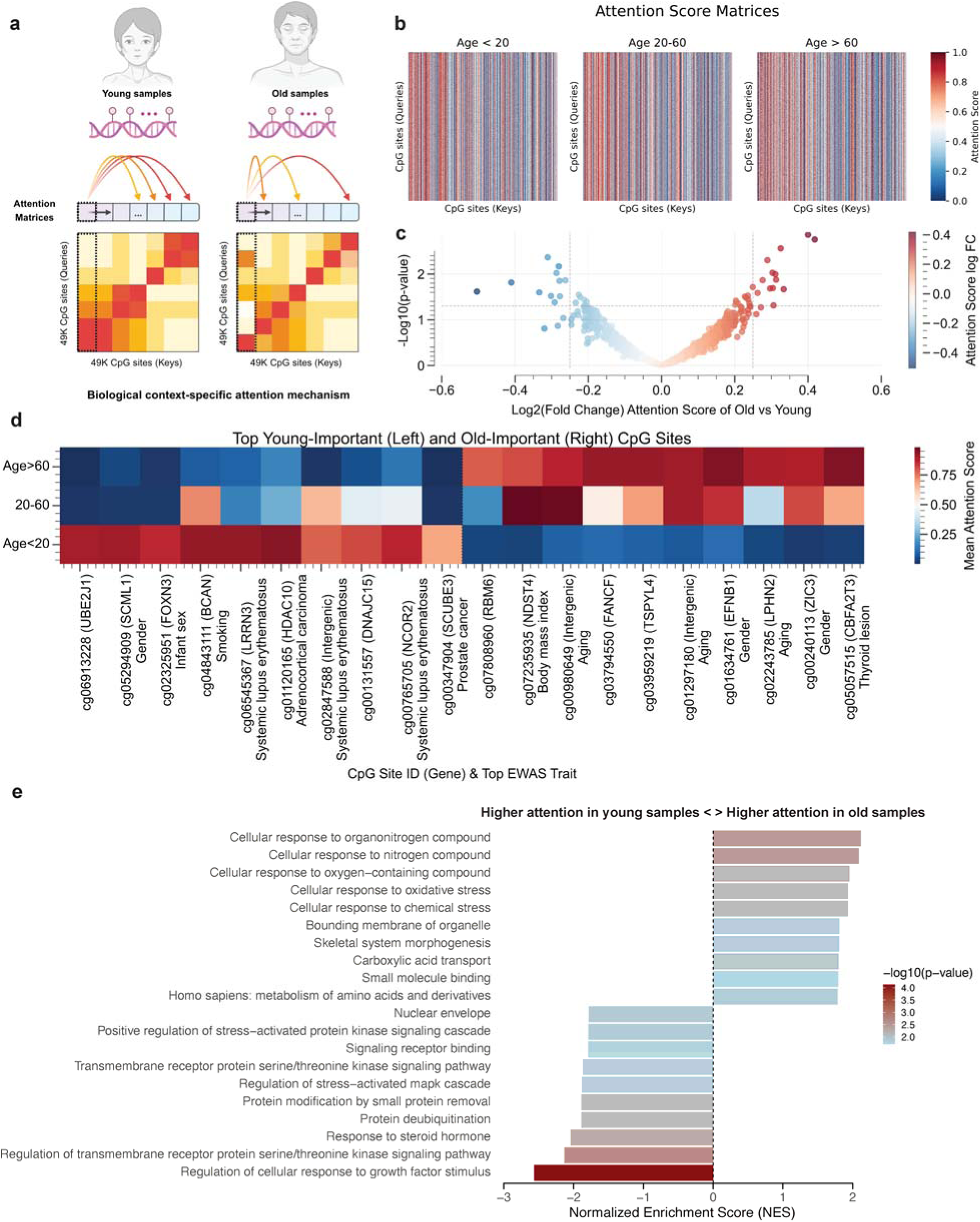
Age-specific attention mechanism analysis. **a.** Schematic comparison of attention patterns between young and old samples, showing differential CpG site interactions. **b.** Attention score matrices across three age groups (<20, 20-60, >60 years), revealing age-specific patterns. **c**. Volcano plot of log p-values versus differential mean attention scores identifies a few influential CpG sites distinguishing the attention pattern of young and old groups. **d.** Heatmap of top young-important (left) and old-important (right) CpG sites, annotated with associated genes and EWAS traits, demonstrating age-specific methylation signatures. **e.** Functional enrichment analysis of top young-important (left) and old-important (right) CpG sites, with bars colored according to -log p-values.

We further analyzed the attention weight distributions across three age groups (< 20, 20-60, and > 60 years) to understand how the model’s attention mechanism adapts to different age ranges (Fig. 5b). The attention patterns revealed systematic shifts in how the model weighs relationships between CpG sites across the lifespan, potentially reflecting underlying biological changes in methylation regulation during aging. Interestingly, attention weights are concentrated on a few CpG sites, suggesting that this sparse set of sites may be significantly relevant to age-specific methylation attention. To identify such statistically influential CpG sites, we extracted sites with large differential attention scores (>1.5 fold change) that were statistically significant (p-value < 0.05) between young and old samples (Fig. 5c). We analyzed the associated EWAS traits and age-specific methylation signatures of the identified important CpGs in both young and old samples (Fig. 5d). In young samples, high-attention CpG sites showed the strongest associations with non-age-associated phenotypes, including sex and autoimmune diseases. Conversely, old samples showed strong attention weights at CpG sites associated with aging, as well as aging-related traits like BMI and thyroid lesions ^35^, validating our model’s biological relevance.

To understand the biological significance of age-specific attention patterns, we performed functional enrichment analysis on CpG sites with differential attention weights between young and old samples. Gene Ontology (GO) and Reactome pathway analysis revealed distinct biological processes associated with high-attention CpG sites in each age group (Fig. 5e). In young samples, highly attended CpG sites were enriched for developmental processes, including cellular response to growth factor stimulus. In contrast, CpG sites receiving higher attention in older samples showed enrichment for oxidative stress and amino acid metabolism. These enrichment patterns validate that MethylGPT’s attention mechanism captures biologically meaningful age-specific methylation signatures.

### Disease risk prediction and intervention analysis

To evaluate MethylGPT’s utility in clinical applications, we analyzed its ability to predict disease risks and assess intervention effects in the Generation Scotland cohort (n = 18,859). We fine-tuned the pre-trained model to predict the risk of 60 major diseases across eight categories, including cardiovascular, respiratory, neurological, and autoimmune conditions, as well as overall mortality, over a 10-year window (Fig. 6a,b). Our results demonstrate that the model achieved an overall Area Under the Curve (AUC) of 0.74 on the validation set and 0.72 on the test set (Fig. 6c).

**Figure 6.**
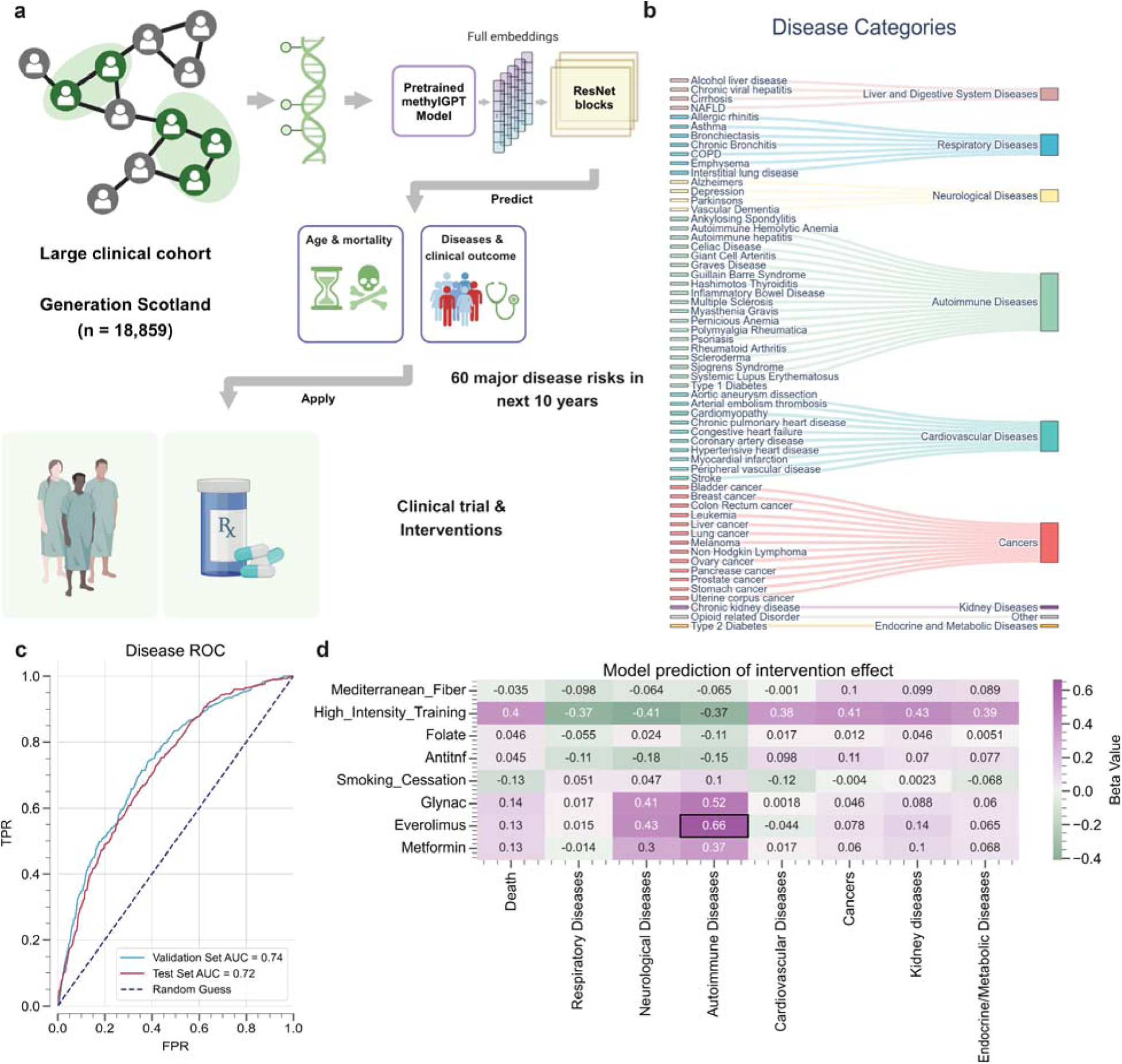
Disease risk prediction and intervention effects using MethylGPT. **a.** Schematic overview of the disease prediction pipeline using Generation Scotland cohort (n = 18,859). The pretrained MethylGPT model processes methylation profiles through ResNet blocks to predict age, mortality, and disease risks, which can then be applied to evaluate clinical interventions. **b.** Visualization of 60 major diseases organized into disease categories (Liver and Digestive System Diseases, Respiratory Diseases, Neurological Diseases, Autoimmune Diseases, Cardiovascular Diseases, Cancers, Kidney Diseases, and Endocrine and Metabolic Diseases). **c.** Receiver Operating Characteristic (ROC) curves showing the overall performance of MethylGPT disease prediction model (seven disease classes and overall mortality) on validation (AUC = 0.736) and test (AUC = 0.720) sets. **d.** Heatmap showing predicted effects (β values) of eight different interventions on disease risks across major disease categories (total n=183): Mediterranean fiber (n=36), high-intensity training (n=5), folate supplementation (n=43), anti-TNF therapy (n=59), smoking cessation (n=16), glyNAC (n=8), everolimus (n=8), and metformin (n=8). Each intervention included an intra-group control as part of the trial design. For phased interventions, only the longest duration timepoint was analyzed. Color scale represents effect size, with purple indicating positive effects (risk reduction) and green indicating negative effects (risk increase). Black box highlights significant effects. Values represent effect size from the Cohen’s d.

Using this disease prediction framework, we systematically evaluated the impact of eight different interventions on predicted disease incidence (Fig. 6d). The model revealed distinct, intervention-specific effects across disease categories. Smoking cessation demonstrated the strongest protective effect against 10-year mortality (β = −0.13) and also reduced cardiovascular disease risk. Notably, high-intensity training showed strong benefits for respiratory, neurological and autoimmune diseases. Similarly, the Mediterranean diet provided modest but consistent protective effects across multiple disease categories, though with varying magnitude.

Interestingly, Everolimus treatment showed a significant risk increase for autoimmune diseases. Although counter-intuitive, this finding is consistent with previous studies showing that prolonged immunosuppressant treatment is associated with an increased incidence of autoimmune diseases ^36^.

Together, these findings demonstrate the potential of MethylGPT for predicting intervention-specific health outcomes and optimizing personalized intervention strategies.

## Discussion

DNA methylation patterns have shown potential as a universal biomarker for disease stratification and monitoring. In oncology, methylation patterns enable the identification of cancer tissue of origin, achieving 81-93% accuracy in predicting primary sites of metastatic tumors and cancers of unknown primary origin ^37^. Methylation-based cardiovascular risk scores demonstrate superior predictive accuracy compared to conventional clinical factors ^38^. Furthermore, methylation markers can predict type 2 diabetes onset years before clinical presentation, providing critical windows for preventive intervention ^39^.

Our results demonstrate that a transformer-based foundation model approach can effectively model DNA methylation patterns while maintaining biological relevance. The organization of CpG sites in the embedding space based on genomic context and regulatory features suggests that MethylGPT captures fundamental aspects of methylation regulation without explicit supervision. This capability addresses a key limitation of traditional linear models that treat CpG sites as independent entities.

The model’s performance in age prediction across diverse tissue types, with improved accuracy over existing methods, demonstrates its potential utility. Particularly notable is the resilience to missing data, maintaining stable performance with up to 70% missingness. This robustness likely stems from the model’s ability to leverage redundant biological signals across multiple CpG sites.

Analysis of age-specific attention patterns revealed distinct methylation signatures between young and old samples. The enrichment of development-related processes in younger samples and aging-associated pathways in older samples, which is consistent with previous studies ^40,41^, suggests that the attention mechanism captures biologically meaningful age-dependent changes in methylation regulation. These findings provide new insights into how methylation patterns evolve across the lifespan.

Several directions for future research emerge from this work. Integration of additional epigenetic features beyond CpG methylation could provide a more comprehensive view of regulatory mechanisms. The development of interpretable attention visualization tools could help bridge the gap between model predictions and biological mechanisms. Additionally, exploring the model’s application to single-cell methylation analysis could reveal cell-type-specific regulatory patterns.

In conclusion, MethylGPT demonstrates how transformer architectures can capture context-dependent methylation patterns while maintaining biological interpretability. The model’s robust performance in handling missing data suggests potential utility in both research and clinical applications.

## Methods

### Pretraining data collection and preprocessing

For the pretraining dataset for the methylGPT, we compose DNA methylation data from 154,063 human samples through the EWAS Data Hub and Clockbase ^28,29^. For quality control, we initially collected data from approximately 300,000 patients and filtered out low-quality entries with high levels of missing data (>40% of total CpG sites). We also applied deduplication to ensure no repetitions in the training data. The cleaned dataset was randomly sampled and quality-checked, covering individuals across 20 distinct tissue types ^42^.

DNA methylation data have varying numbers of CpG entries depending on the array platform (Illumina 27k, Illumina 450k, and EPIC). To address these differences and ensure biological relevance, we focused on 49,156 CpG sites selected based on importance by EWAS traits ^30^ and array format compatibility. In detail, these 49,156 CpG sites satisfy either (1) CpG are associated with more than 5 traits according to EWAS catalog or (2) CpG appears in more than 95% of the pretraining dataset. All methylation values were normalized using standard protocols. Missing values were marked for downstream masked prediction tasks.

Data is processed into a matrix of *x* ɛ ℝ*^N^*^×*M*^, where each element *x_i,j_* denotes the magnitude of methylation of a CpG site *j* in sample *i*. *N* is the number of samples and *M* is the number of CpG sites (i.e. 49,156).

### Model architecture

MethylGPT consists of three main components: an embedding module, a transformer module, and task-specific heads. The input data *x* is tokenized and fed into the modules consecutively.

We depict the input tokenization and the module details as follows.

### CpG site tokenization

The processed data contains methylation readings of *M* (49,156) CpG sites. For each site *c_ij_* (*j* ɛ {0,1,…, *M*}), we assign an integer identifier *id*(*c_ij_*). The full CpG tokens for an individual sample are 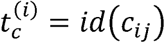.

### Embedding layers

We utilize the embedding layers for the CpG site tokens to map each token to a fixed-length embedding vector of dimension D. We employ fully connected layers for the methylation values to encode the methylation level into vector embeddings and maintain the ordinal relation of the values.

For each CpG site, the embedding module projects both CpG site identifiers and their methylation values into separate embeddings (referred to as CpG embeddings and methylation value embeddings), which are then merged through an element-wise sum. The final embedding for sample i is defined as:

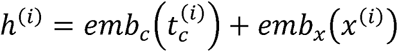

The embedding dimension is set to 64. A special [CLS] token is prepended to each sequence for learning sample-level representations.

### Transformer module

We employ the self-attention transformer ^17,43^ to encode the complete input embedding. The transformer module comprises 6 transformer blocks, each containing a multi-head self-attention layer (4 heads) and a standard MLP layer. Layer normalization and residual skip connections are applied after each layer. The self-attention mechanism operates on the sequence of M embedding vectors, making it particularly suitable for capturing interactions between CpG sites. The transformer processes the sequence according to:

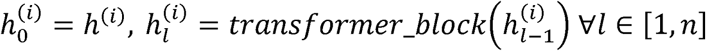

We utilize the resulting representation 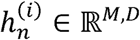 for both CpG-level and sample-level tasks. The self-attention layer leverages FlashAttention for efficient training and inference ^44^. The model dimension is set to 64, with also an intermediate dimension of 64 in the feed-forward layers. The transformer module processes a sequence of input embeddings comprising 49,157 sites with 64 dimensions and outputs “contextualized embeddings” of the same shape.

The input dimension M can reach tens of thousands of CpG sites, consuming huge memory and creating a significant challenge for efficient model training. We leverage the Flash-Attention ^44^ implementation as a tool to greatly accelerate the training and inference of the model while minimizing memory footprint.

The task-specific heads attached to the transformer process contextualized embeddings into diverse predictions specific to the task. In the pre-training phase, a linear layer projects output embeddings of each CpG site to predict the methylation value. In the fine-tuning phase, the MLP or convolutional layers process the complete output embeddings to predict biological age or occurrence of disease.

### Model pretraining

The model was trained on two complementary objectives. First, we randomly masked 30% of CpG sites (i.e., their value embeddings were excluded from the input embedding process) and trained the model to reduce the MSE between the predicted and original methylation values at the masked CpG sites. The Methylation Value Prediction (MVP) objective is defined as:

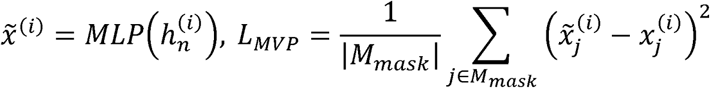

where 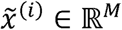 represents the row of predicted methylation value estimates for sample i. The MVP objective encourages the model to effectively encode relationships among the CpG sites in the dataset.

Second, a profile reconstruction task used the [CLS] token output embedding to reconstruct complete methylation profiles, as also described in a previous study ^25^. The model feeds the [CLS] token’s output embedding from the previous step back into the [CLS] token input, while all other tokens are masked. The objective of the profile reconstruction task is:

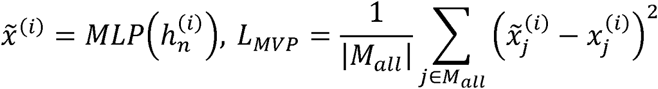

Training was performed using the AdamW optimizer with a learning rate of 0.001, The model was trained for 10 epochs with a batch size of 16 on NVIDIA A100 GPU. The learning rate is set to decay 10% after each epoch.

### Evaluation metrics

Model performance was assessed through multiple metrics. We calculated the MSEand MAE for methylation value prediction, along with Pearson correlation coefficients between predicted and actual methylation values. For age prediction tasks, we measured accuracy using MedAE in years. For disease prediction tasks, model effectiveness was evaluated using the AUC, which measures the model’s discriminative ability to differentiate between various disease states.

### Age prediction experiments

For the evaluation of age prediction, we utilized a dataset comprising 13,505 samples with 21,368 CpG sites ^31^. From the accompanying metadata, we identified training (5,461 samples), validation (1,366 samples), and test (4,626 samples) sets, with a total of 49,156 CpG sites.

We fine-tuned pre-trained MethylGPT using the downstream prediction head ResNet1D. The ResNet1D consists of six residual blocks, where each residual block includes two 1D convolutional layers with a kernel size of 3, each followed by batch normalization and ReLU activation. Specifically, we input 49,156 CpG sites into the MethylGPT, generating an embedding with dimensions (49,156, 64). To reduce dimensionality, this embedding was passed through a 3x3 1D convolutional layer, condensing the feature space to 32 channels. The reduced-dimensionality output was subsequently fed into six residual blocks, followed by an average pooling layer and a linear layer for age prediction. Both the pre-trained MethylGPT and the downstream ResNet1D prediction head were trained using the MSE loss function as the optimization objective.

To assess robustness, we systematically masked increasing proportions (10-90%) of CpG sites in the test set and evaluated prediction performance. Comparison methods (ElasticNet, MLP, Horvath’s clock) were trained and evaluated on the same data splits.

### Disease prediction and intervention evaluation

We fine-tuned the pre-trained model, maintaining consistency with the downstream prediction head architecture, ResNet1D, used in age prediction to demonstrate the generalizability of the pre-trained model across downstream tasks. By utilizing the same downstream network structure for both age and disease prediction, we aimed to confirm that the model’s effectiveness was not due to meticulous architecture optimization but rather due to its inherent flexibility.

To evaluate this, we curated datasets from the Generation Scotland cohort (n = 18,859), comprising 1,378 samples for training, 295 for validation, and 296 for testing. In fine-tuning, the model was trained to simultaneously predict the risk of 60 major diseases across eight categories, leading to the development of a comprehensive disease prediction model. For each disease category, a sample was labeled as ‘1’ if the disease was present and ‘0’ otherwise. This multi-label classification task, where a sample could have one or multiple co-occurring diseases, introduced substantial complexity to the prediction challenge. Both the pre-trained MethylGPT and the downstream ResNet1D prediction head were optimized using the cross-entropy loss function.

To further explore the impact of interventions on predictive outcomes, we applied the disease prediction model to assess data from eight types of interventions across six GEO datasets (GSE219217 ^45^, GSE268211 ^46^, GSE176325 ^47^, GSE191297 ^48^, GSE201532 ^49^, GSE276988 ^50^), encompassing a total of 183 samples. The interventions examined in this study included Mediterranean fiber (n=36), high-intensity training (n=5), folate supplementation (n=43), anti-TNF therapy (n=59), smoking cessation (n=16), glyNAC (n=8), everolimus (n=8), and metformin (n=8). Each intervention included an intra-group control as part of the trial design. For the phased interventions, only the longest duration of each intervention was retained for analysis.

### Attention analysis

Age-specific attention patterns were analyzed by extracting attention scores from all heads in the final transformer layer. We computed mean attention scores for each CpG site across samples within defined age groups (<20, 20-60, >60 years). CpG sites with significantly different attention scores between age groups were identified using two-sided t-tests with Benjamini-Hochberg correction.

For CpG sites showing differential attention patterns between age groups, we performed Gene Ontology (GO) and Reactome gene set enrichment analysis using MethylGSA ^51^.

### Statistical analysis

All statistical tests were two-sided unless otherwise specified. Error bars in the figures represent standard deviation across samples. Sample sizes and statistical methods are specified in figure legends.

## Code availability

The MethylGPT code and pre-trained models will be made available on github upon publication.

## Data availability

All methylation data used in this study are available through EWAS Data Hub, GEO, and Clockbase. Processed datasets and analysis scripts will be deposited to github upon publication.

## Acknowledgments

This work was supported by grants from the National Institute on Aging and Hevolution to VNG. It was also supported by the James Fickel Foundation. KY was supported by National Institute on Aging grant F99AG088431. We thank the Biomarkers of Aging Consortium and Generation Scotland for providing access to their datasets. Generation Scotland received core support from the Chief Scientist Office of the Scottish Government Health Directorates [CZD/16/6] and the Scottish Funding Council [HR03006] and is currently supported by the Wellcome Trust [216767/Z/19/Z]. Genotyping of the Generation Scotland samples was carried out by the Genetics Core Laboratory at the Edinburgh Clinical Research Facility, University of Edinburgh, Scotland and was funded by the Medical Research Council UK and the Wellcome Trust (Wellcome Trust Strategic Award “STratifying Resilience and Depression Longitudinally” (STRADL) Reference 104036/Z/14/Z). The DNA methylation profiling and analysis was supported by Wellcome Investigator Award 220857/Z/20/Z and Grant 104036/Z/14/Z (PI: Prof AM McIntosh) and through funding from NARSAD (Ref: 27404; awardee: Dr DM Howard) and the Royal College of Physicians of Edinburgh (Sim Fellowship; Awardee: Prof HC Whalley)

## Author contributions

KY conceived the idea and designed the study. KY, SL, and HL collected initial data. KY, JS, and HC designed the model and performed pre-training. KY and YZ performed model finetuning and analysis. XC and AE helped with data analysis. DLM, REM, and MM helped with human cohort data curation. KY, JS, and HC wrote the manuscript. All authors edited and contributed to the manuscript. BW and VNG supervised the study.

